# Micro-C-TALE: BAC-based targeted high-resolution measurement of 3D chromatin interactions

**DOI:** 10.64898/2026.10.21.760768

**Authors:** Anna Golberg, Arkadiy K Golov, Revital Shemer, Noam Kaplan

**Author notes:** equal contribution (Anna Golberg, Arkadiy K Golov).

## Abstract

Chromosome conformation capture methods have greatly advanced our understanding of genome organization, but obtaining high-resolution interaction maps of selected genomic regions remains costly due to the sequencing requirements. Here, we introduce Micro-C-TALE, a new chromosome conformation capture method that combines Micro-C, which uses micrococcal nuclease to improve the resolution and sensitivity of chromatin interaction measurements, and C-TALE, which uses probes generated from bacterial artificial chromosomes (BACs) for targeted enrichment. This combination enables the generation of high-resolution interaction maps without requiring custom synthesis of capture oligonucleotides. We applied Micro-C-TALE to a 214 kb region in human HAP1 cells, obtaining detailed interaction maps with approximately 5-6 million sequenced read pairs. Compared with published genome-wide Micro-C data, Micro-C-TALE yielded a higher amount of informative interactions within the targeted region despite ∼550-fold lower sequencing depth. To evaluate the sensitivity of the method to fine-scale changes in genome organization, we introduced a heterozygous CTCF binding site into the targeted region using CRISPR-Cas9. Comparing Micro-C-TALE to the restriction enzyme-based C-TALE, we found Micro-C-TALE yielded a ∼5-fold higher fraction of informative reads and detected novel CTCF-mediated point interactions associated with the inserted CTCF site, which were barely discernible using C-TALE. Our results demonstrate that Micro-C-TALE provides a flexible and cost-effective approach for targeted high-resolution measurement of chromatin interactions, facilitating the investigation of fine-scale genome organization and its response to genetic perturbations.

## Introduction

The spatial organization of the genome is closely linked to various nuclear processes, including transcription, DNA replication, DNA repair and cell division^1–3^. Over the past two decades, chromosome conformation capture (3C)-based methods have greatly advanced our understanding of genome organization by allowing high-throughput measurement of spatial interactions between genomic loci^4–7^, and have also been leveraged for genome assembly applications^8–11^. In particular, Hi-C enables genome-wide measurement of pairwise chromatin interactions, revealing organizational features at different genomic scales, including genomic compartments, Topologically Associating Domains (TADs), and point interactions^5,6,12–15^. Some of these structural features have been linked to specific molecular mechanisms, such as the role of cohesin-mediated loop extrusion and CTCF in the formation of TADs and point interactions in vertebrates^16–19^. However, despite important methodological improvements, obtaining cost-effective highly-detailed chromatin interaction maps remains challenging, particularly when studying fine-scale interactions between individual genomic elements.

The resolution of Hi-C interaction maps is limited both by the size of restriction fragments and by sequencing depth^6,7^. Thus, obtaining detailed interaction maps of specific genomic loci in mammalian genomes may require extensive genome-wide sequencing, making such experiments costly and inefficient. To address this limitation, several targeted chromosome conformation capture methods have been developed, allowing deeper sampling of selected genomic regions while reducing sequencing requirements. Early approaches such as 4C and 5C enabled high-throughput measurement of interactions involving selected viewpoints or multiple predefined genomic fragments^20–22^. Subsequently, hybridization-based enrichment approaches were developed, using biotinylated probes to capture selected proximity-ligation products. These include Capture-C, NG Capture-C, Capture Hi-C and promoter Capture Hi-C, which enable detailed analysis of interactions involving selected genomic viewpoints, such as gene promoters^23–26^. Other approaches, including Targeted Chromatin Capture (T2C), Hi-C² and Tiled-C, use capture probes spanning extended genomic regions to generate high-resolution interaction maps^27–29^. C-TALE (Chromatin TArget Ligation Enrichment) provides an alternative regional targeting strategy using probes generated from bacterial artificial chromosomes (BACs), avoiding the need for custom oligonucleotide synthesis and providing a flexible and cost-effective approach for targeted enrichment^30^.

An important advance in chromosome conformation capture was the development of Micro-C, which substitutes restriction enzyme digestion with micrococcal nuclease (MNase) digestion^31–34^. MNase preferentially digests linker DNA between nucleosomes, allowing chromatin fragmentation at approximately nucleosome-scale resolution. Unlike restriction enzyme-based approaches, MNase digestion is not constrained by the distribution of specific recognition sequences, enabling the measurement of chromatin interactions at substantially finer genomic scales. Initially developed in yeast, Micro-C was subsequently adapted to mammalian genomes, revealing fine-scale chromatin organization and point interactions that are difficult to resolve using conventional Hi-C^33,34^. Importantly, beyond its improved fragmentation resolution, Micro-C and some of its variants have been shown to provide a higher signal-to-noise ratio and increased sensitivity to point interactions compared with Hi-C, enabling significantly improved detection of interactions involving CTCF binding sites, enhancers and promoters^33–36^. These capabilities have facilitated high-resolution investigation of enhancer-promoter communication and the relationship between transcriptional regulation and chromatin interactions^36,37^. However, achieving these higher resolutions still requires massive sequencing depth, as the space of possible interactions grows quadratically with the number of interrogated loci. Thus, obtaining sufficiently sampled interaction maps at nucleosome-scale resolution remains challenging, particularly when studying individual genomic regions within large mammalian genomes.

To overcome the sequencing depth limitations of genome-wide Micro-C, several approaches have recently been developed combining MNase-based fragmentation with targeted enrichment of chromatin interactions. Micro-Capture-C (MCC) uses hybridization-based capture of selected genomic viewpoints, enabling the measurement of interactions between regulatory elements at near base-pair resolution^38,39^. Tiled Micro-Capture-C (TMCC) extends this approach by using densely tiled capture probes across extended genomic regions, allowing the generation of high-resolution contact maps and the investigation of fine-scale chromatin structures and their dependence on cohesin and CTCF^40^. Similarly, Region Capture Micro-C (RCMC) combines Micro-C with targeted enrichment of extended genomic regions, enabling exceptionally deep sequencing and revealing previously unresolved interactions between enhancers and promoters, including highly nested structures termed microcompartments^41^. Other MNase-based approaches, including MChIP-C and Micro-C-ChIP, use chromatin immunoprecipitation to enrich interactions associated with specific chromatin proteins or histone modifications, enabling high-resolution characterization of chromatin interactions without restricting the analysis to predefined genomic regions^36,42^. Together, these methods demonstrate the utility of combining MNase-based fragmentation with targeted enrichment for studying fine-scale genome organization. However, methods based on hybridization capture of extended genomic regions generally require custom-designed and synthesized oligonucleotide probes, motivating the development of alternative targeting strategies.

Here, we present Micro-C-TALE, a targeted chromosome conformation capture method combining the nucleosome-scale fragmentation of Micro-C with the cost-effective BAC-based enrichment strategy of C-TALE. We apply Micro-C-TALE to a selected genomic region in human HAP1 cells and characterize the resulting interaction maps and sequencing efficiency. Finally, to evaluate the ability of Micro-C-TALE to detect fine-scale changes in chromatin organization, we use CRISPR-Cas9 to introduce a CTCF binding site into the targeted region and compare the resulting interaction maps with those obtained using C-TALE. Our results demonstrate the applicability of BAC-based enrichment to Micro-C and provide an alternative approach for high-resolution analysis of chromatin interactions in selected genomic regions.

## Results

### Overview of Micro-C-TALE

To enable high-resolution measurement of chromatin interactions in selected genomic regions, we developed Micro-C-TALE, combining the MNase-based chromatin fragmentation of Micro-C with the BAC-based targeting strategy of C-TALE (**Figure 1**; step-by-step protocol provided in Supplementary Methods). The protocol consists of three main stages: Micro-C library preparation, production of BAC-derived capture probes, and targeted enrichment of Micro-C libraries.

**Figure 1.**
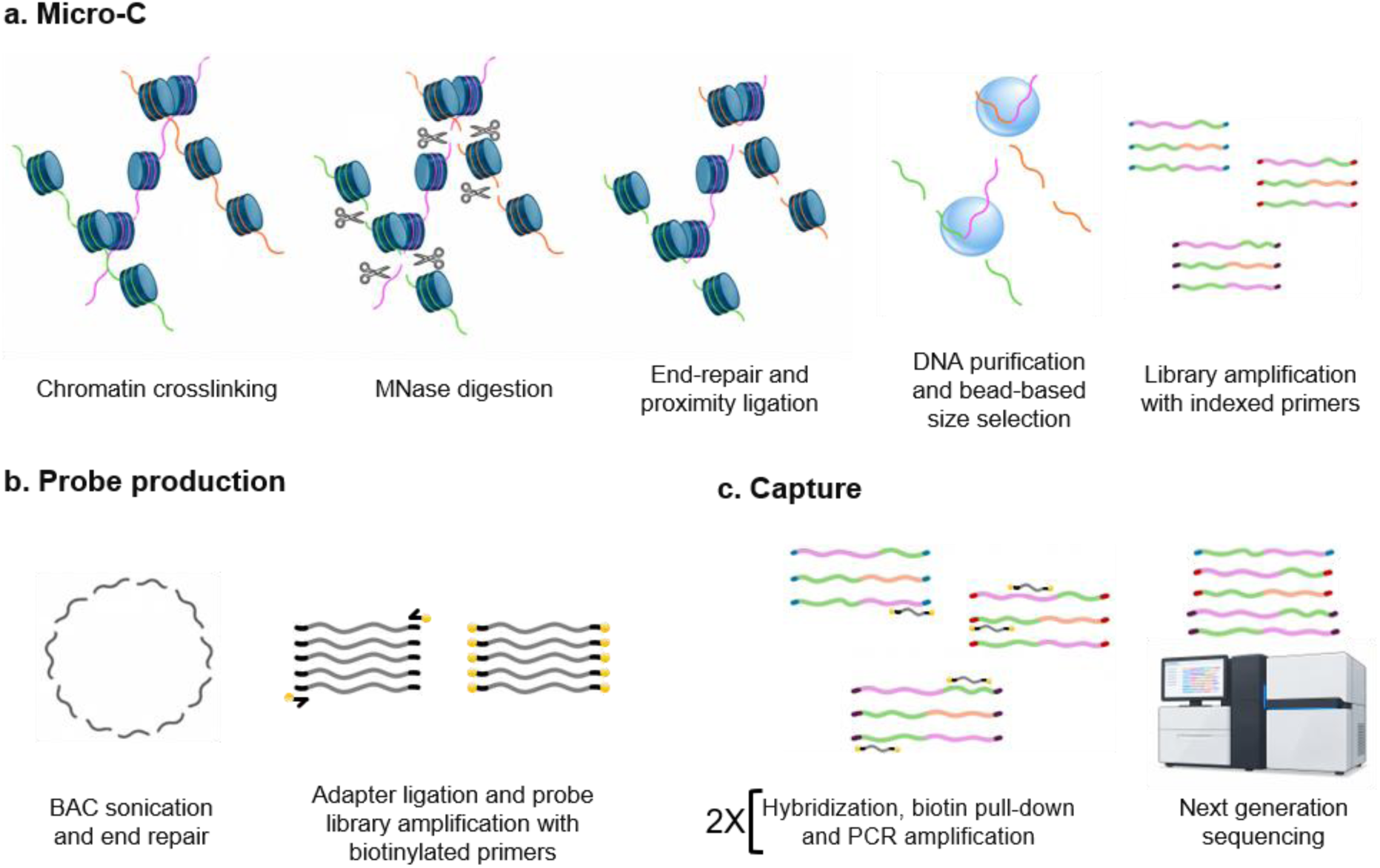
Overview of Micro-C-TALE. Schematic representation of the three main stages of the Micro-C-TALE protocol. **(a)** Micro-C library preparation, including chromatin crosslinking, MNase digestion, end repair and proximity ligation, DNA purification and bead-based size selection, and library amplification using indexed primers. **(b)** Production of biotinylated capture probes from bacterial artificial chromosome (BAC) DNA by sonication, end repair, adapter ligation and PCR amplification using biotinylated primers. **(c)** Two rounds of targeted enrichment by hybridization with BAC-derived probes, streptavidin pull-down and PCR amplification, followed by next-generation sequencing.

For Micro-C library preparation, cells are first crosslinked with formaldehyde, followed by cell lysis and chromatin digestion using micrococcal nuclease (MNase). The amount of MNase is calibrated to obtain predominantly mononucleosomal fragments while avoiding excessive digestion. Following MNase digestion, DNA ends are repaired and subjected to proximity ligation. In contrast to conventional Micro-C, we omit biotin labeling and streptavidin pull-down of ligation junctions, aiming to minimize potential loss of library complexity, as previously suggested for C-TALE^30^. Following reversal of crosslinking and DNA purification, we use AMPURE XP bead-based size selection to enrich for ligated fragments and remove unligated mononucleosomal DNA. The resulting DNA is converted into an indexed Illumina sequencing library.

In parallel, BAC-derived capture probes are prepared according to the general strategy of C-TALE^30^. BAC clones covering the genomic region of interest are isolated and fragmented by sonication. The fragments are ligated to custom adapters and amplified using biotinylated primers, producing a pool of double-stranded DNA probes covering the targeted genomic region. This strategy avoids the need for custom synthesis of capture oligonucleotides and allows the production of additional probes by PCR amplification.

Finally, the Micro-C libraries are subjected to two rounds of hybridization-based enrichment using the BAC-derived probes. Following each round of hybridization, probe-bound DNA fragments are recovered using streptavidin beads, washed and PCR-amplified. The resulting enriched libraries are sequenced to produce high-resolution interaction maps of the targeted genomic region.

### Generation of a targeted high-resolution interaction map with Micro-C-TALE

To evaluate the performance of Micro-C-TALE, we applied the method to human HAP1 cells, targeting a 214 kb region on chromosome 1 (Chr1:30,754,221–30,968,290, hg38) using BAC-derived capture probes. We generated Micro-C libraries from wild-type HAP1 cells, followed by two rounds of targeted enrichment and paired-end sequencing.

We obtained approximately 5.8 million sequenced read pairs, of which ∼2.3 million represented uniquely mapped interactions. Of these, approximately 873,000 unique read pairs mapped on both sides to the targeted region, corresponding to 38% of all unique read pairs. As expected for MNase-based chromatin fragmentation without biotin enrichment of ligation fragments, a large proportion of these interactions occurred at short genomic distances. Restricting the analysis to interactions spanning at least 1 kb or 5 kb, we obtained approximately 107,000 and 60,000 unique on-target interactions, respectively. These favorably compare to a recent genome-wide Micro-C experiment in which ∼3,186 million read pairs were sequenced (550x more than Micro-C-TALE), resulting in a lower amount of informative reads for the same target region (∼82,000 unique on target >1kb; ∼26,000 unique on target >5kb). Thus, despite the relatively modest sequencing depth, these results indicate substantial enrichment of interactions involving the targeted region.

We next examined the resulting interaction map at 2Kb resolution and compared it with a HAP1 Hi-C map^18^ of the same region (**Figure 2**). While the Hi-C map appears extremely noisy at this resolution, the Micro-C-TALE map shows clearly outlined structures including TADs and point interactions. Specifically, CTCF-CTCF point interactions anchored at the TAD boundaries are clearly observable and accurately positioned in the Micro-C-TALE map, but barely observable in the Hi-C map. These results demonstrate that Micro-C-TALE can generate targeted chromatin interaction maps from relatively modest sequencing depths, allowing the detection of fine-scale features of genomic organization that may be missed by standard Hi-C.

**Figure 2.**
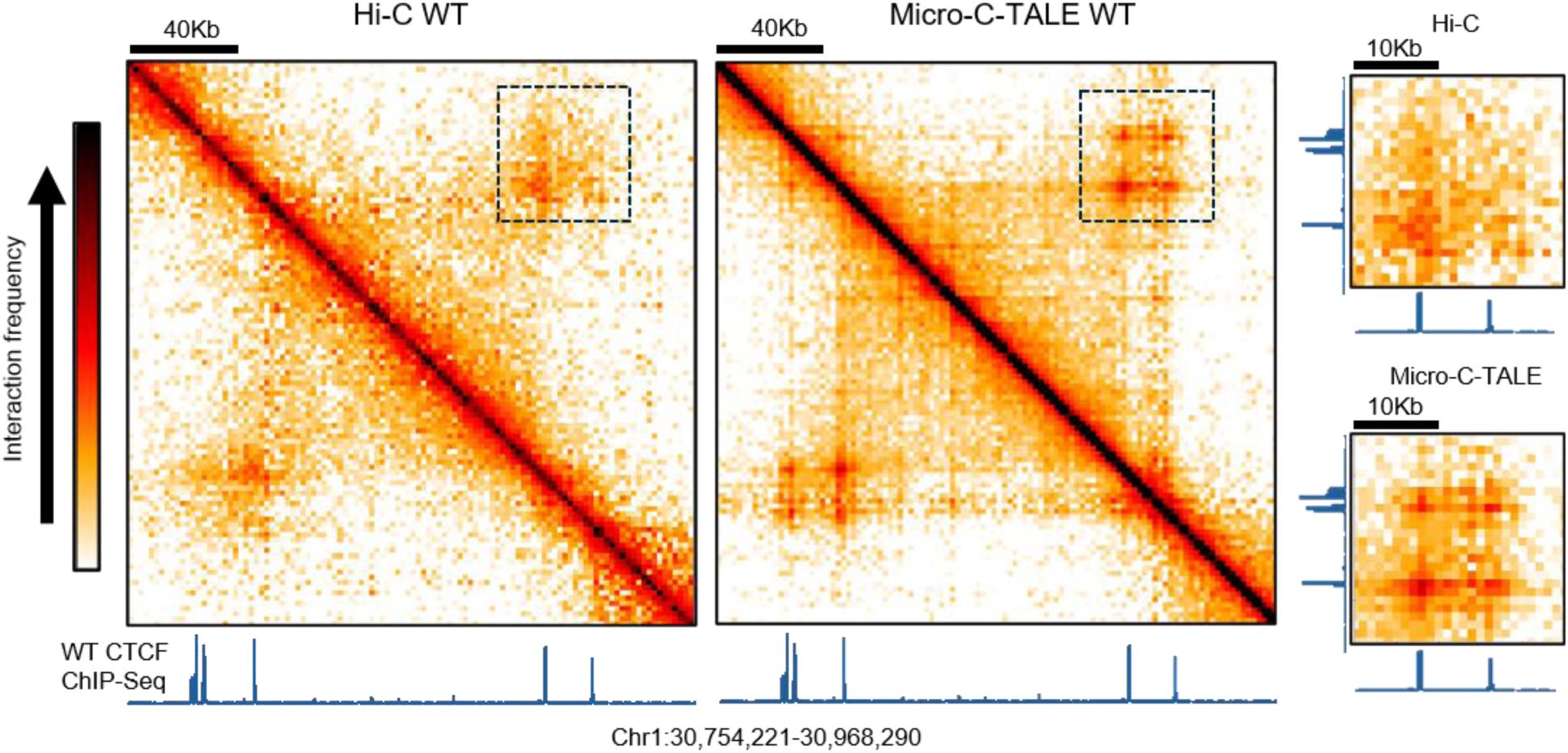
Comparison of Micro-C-TALE and Hi-C interaction maps. Micro-C-TALE and Hi-C 2kb resolution interaction maps of target region in HAP1 cells. Hi-C was taken from Sanborn et al.^18^. HAP1 wildtype CTCF ChIP-Seq from Haarhuis et al. ^43^ is shown below in blue. Dashed squares indicate regions which are enlarged on the right, containing CTCF-associated point interactions.

### Micro-C-TALE detects point interactions induced by heterozygous CTCF binding site insertion

Next, we asked whether Micro-C-TALE is sensitive enough to detect subtle small-scale changes in 3D genome organization. To this end, we used CRISPR-Cas9 to introduce a single CTCF binding site into the target region. Sanger sequencing showed a combination of the wildtype and mutant alleles, suggesting that the HAP1 cells underwent diploidization prior to the insertion^44^. We reasoned that the insertion of a single heterozygous CTCF binding site might create CTCF-CTCF point interactions that would require a highly-sensitive assay to detect. We performed Micro-C-TALE on the heterozygous CTCF insertion clone, resulting in 5.3 million read pairs which yielded ∼85,000 and ∼47,000 uniquely mapped on-target reads with genomic distances >1kb and >5kb respectively. We then compared the interaction maps visually (**Figure 3a**). Both maps showed TAD structures and point interactions. Reassuringly, the interaction map of the CTCF insertion clone showed new point interactions emerging at the intersection of the CTCF insertion site with other established CTCF interacting sites. To further examine this change, we used a virtual 4C to plot the interactions of the target insertion locus with its downstream genomic vicinity. We observe the appearance of two interacting peaks at the locations of known HAP1 CTCF sites, consistent with the formation of new CTCF-CTCF point interactions (**Figure 3b**). Importantly, these new interactions are apparent despite their signal intensity being reduced by approximately half due to heterozygosity (effectively averaging with the non-interacting wildtype allele). These results indicate that Micro-C-TALE can be sensitive enough to detect subtle structural changes in localized regulatory genomic features.

**Figure 3.**
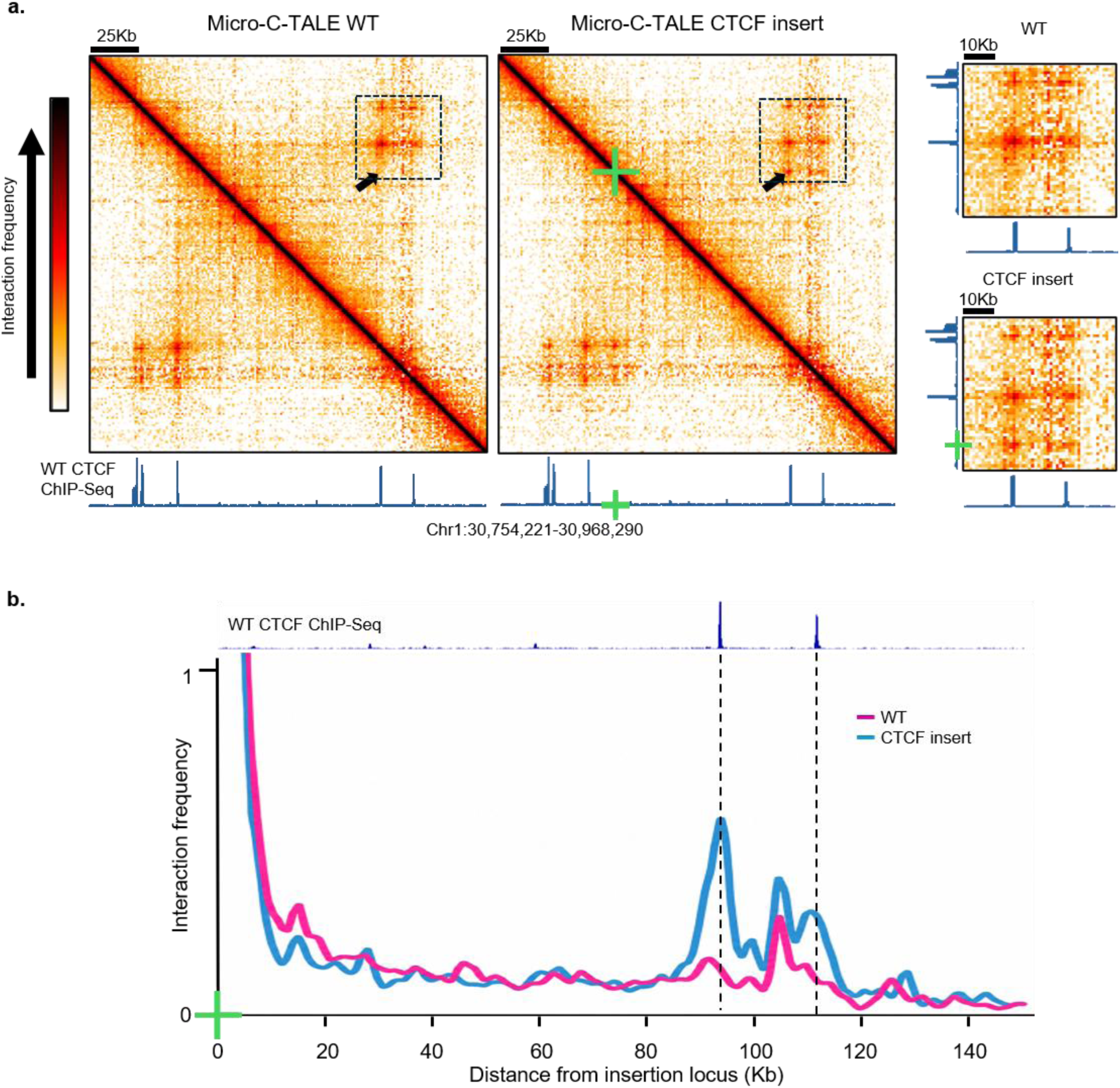
Detection of CTCF-associated point interactions following insertion of a CTCF binding site. **(a)** Micro-C-TALE 1kb resolution interaction maps of target region for wildtype and CTCF insert clone. HAP1 wildtype CTCF ChIP-Seq from Haarhuis et al. ^43^ is shown in blue below and on the side of interaction maps. Dashed squares indicate regions which are enlarged on the right. The green cross indicates the CTCF insertion site, and the arrow indicates an emerging CTCF-associated point interaction. **(b)** Virtual 4C plot showing the interactions of the insertion locus (Chr1:30,817,713) with its downstream genomic vicinity. Wildtype is shown in magenta and the CTCF insertion clone in blue. HAP1 wildtype CTCF ChIP-Seq is shown above.

### Micro-C-TALE improves on C-TALE resolution, sensitivity and efficiency

To further dissect whether the improved interaction map quality of Micro-C-TALE is due to sequencing depth or MNase usage, we generated a C-TALE interaction map for the same target region, using the same HAP1 CTCF-insertion clone. We sequenced ∼16 million read pairs from the C-TALE library, ultimately yielding ∼49,000 and ∼36,000 uniquely mapped on-target reads with genomic distances >1kb and >5kb respectively. Comparatively, this gives 0.3%/0.2% (>1kb/>5kb) uniquely mapped on-target reads out of total sequenced reads for C-TALE, relative to 1.6%/0.9% and 1.9%/1.0% for the Micro-C-TALE libraries. This suggests that Micro-C-TALE is also more efficient than C-TALE in capturing informative DNA fragments, in spite of these protocols sharing the probe preparation and capture steps. We then compared the C-TALE and Micro-C-TALE interaction maps of the HAP1 CTCF-insertion clone (**Figure 4**). Interestingly, despite its targeted nature, the C-TALE map did not show obvious CTCF-CTCF point interactions, consistent with previous observations regarding the improved sensitivity of MNase-based assay in detecting such interactions. Thus, our results suggest that Micro-C-TALE may provide higher resolution, better sensitivity and increased efficiency than C-TALE.

**Figure 4.**
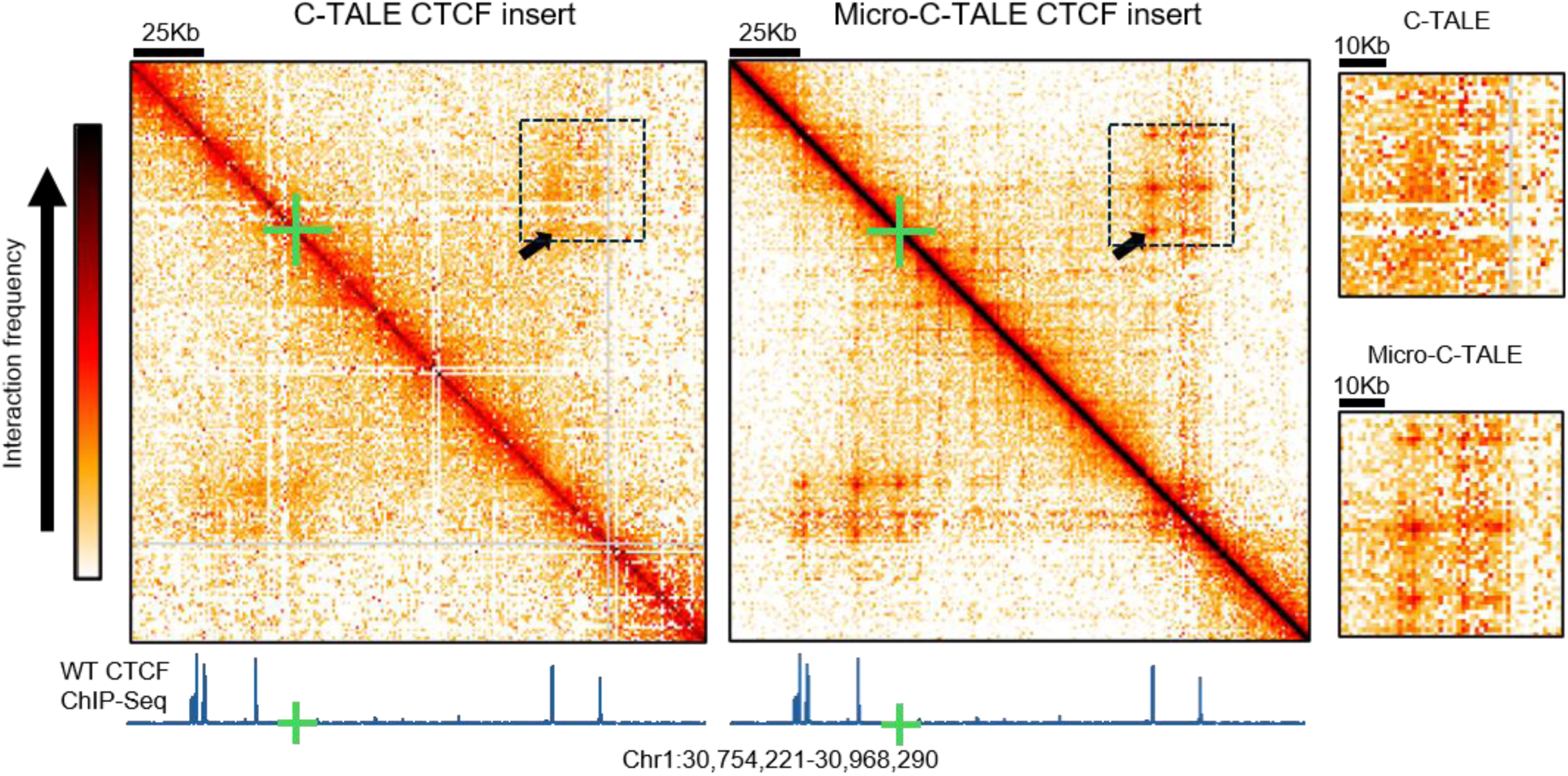
Comparison of Micro-C-TALE and C-TALE interaction maps. Interaction frequency maps of the targeted region in the HAP1 CTCF insertion clone, generated using Micro-C-TALE and C-TALE at 1 kb resolution. HAP1 wildtype CTCF ChIP-Seq from Haarhuis et al. ^43^ is shown below in blue. Dashed squares indicate regions which are enlarged on the right. The green cross indicates the CTCF insertion site, and the arrow indicates a CTCF-associated point interaction that is clearly detected by Micro-C-TALE but is barely discernible by C-TALE.

## Discussion

In this work we developed Micro-C-TALE, a targeted chromosome conformation capture method combining the nucleosome-scale fragmentation of Micro-C with the BAC-based enrichment strategy of C-TALE. We demonstrate that Micro-C-TALE enables the generation of detailed chromatin interaction maps of selected genomic regions of interest using modest sequencing depth.

An important advantage of Micro-C-TALE is the use of BAC-derived probes for targeted enrichment. In contrast to methods relying on custom-designed oligonucleotide panels, BAC-based probes can be generated using standard molecular biology procedures without requiring custom oligonucleotide synthesis. Once prepared, the BAC-derived probe libraries can be amplified to produce additional material for subsequent experiments. This provides a flexible and cost-effective approach, particularly for studies focusing on extended genomic regions or requiring comparisons across multiple experimental conditions. Micro-C-TALE thus complements existing targeted Micro-C approaches, including MCC, TMCC and RCMC, by providing an alternative strategy for targeted enrichment.

The increased fragmentation resolution and sensitivity of Micro-C-TALE provide important advantages over restriction enzyme-based approaches such as Hi-C and C-TALE. Unlike restriction enzyme digestion, MNase digestion is not constrained by the distribution of specific recognition sequences, enabling more detailed characterization of fine-scale chromatin interactions, including point interactions involving regulatory elements. In addition, previous studies have demonstrated that Micro-C provides increased sensitivity to point interactions compared with Hi-C^33,34,36,37^. We demonstrate this advantage experimentally by introducing a CTCF binding site into the targeted genomic region and directly comparing the resulting interaction maps generated using Micro-C-TALE and C-TALE. The point interaction associated with the inserted CTCF site is clearly detected using Micro-C-TALE - despite it being heterozygous - but is barely discernible using C-TALE, demonstrating the improved sensitivity of Micro-C-TALE to fine-scale changes in chromatin organization.

Another characteristic of Micro-C-TALE is the omission of biotin labeling and streptavidin pull-down of proximity-ligation junctions during library preparation. This choice was motivated by the possibility of preserving library complexity, as previously suggested for C-TALE^30^. However, omitting this junction pull-down step also results in the retention of DNA fragments that do not contain ligation junctions, potentially reducing the fraction of informative sequencing reads. We partially address this limitation by size selection to enrich for ligated fragments. Since targeted enrichment substantially reduces the sequencing depth required to obtain detailed maps of selected regions, this trade-off is often acceptable in applications where library complexity is particularly important. Nevertheless, further optimization and direct comparisons would be necessary to determine the effects of this choice on library complexity and sequencing efficiency.

Micro-C-TALE retains some limitations inherent to hybridization-based capture, including variation in enrichment efficiency across genomic regions and dependence on the availability and coverage of suitable BAC clones. Furthermore, the relatively extensive hybridization and enrichment procedure adds experimental time and complexity compared with standard genome-wide approaches. As with other chromosome conformation capture methods, the resulting interaction maps represent population-averaged interaction frequencies rather than direct measurements of three-dimensional distances.

In summary, Micro-C-TALE provides a cost-effective strategy for targeted, high-resolution measurement of chromatin interactions. By combining MNase-based fragmentation with BAC-derived regional enrichment, the method enables detailed investigation of chromatin organization in selected genomic regions and facilitates the detection of fine-scale structural changes following genetic or other experimental perturbations.

## Methods

Complete step-by-step Micro-C-TALE protocols are provided in the Supplementary Methods.

### Cell culture and fixation

HAP1 cells were cultured in IMDM medium supplemented with 10% fetal bovine serum and 1% penicillin–streptomycin. Cells were harvested using trypsin, counted, and centrifuged at 500 × g for 5 min at room temperature. Cells were resuspended in 950 µl PBS and 55 µl 37% Formaldehyde for every 10⁷ cells. The cells were then crosslinked for 10 min at room temperature with gentle agitation. Crosslinking was quenched by dropwise addition of 600 µl of 2 M Tris-HCl (pH 7.4) per 10⁷ cells, followed by 5 min of rocking and 10 min on ice. Cells were centrifuged at 1,000 × g for 5 min at 4 °C, washed with PBS containing 0.1 M Tris-HCl, and divided into aliquots of 5 × 10⁶ cells. Following centrifugation, cell pellets were snap-frozen in liquid nitrogen and stored at −80 °C.

### BAC-based probe production

A total of 1.3 µg of BAC DNA (CH17-219E19; chr1:30,754,221–30,968,290) was fragmented by sonication using a Covaris system to obtain fragments of 200–300 bp. Fragmented DNA was subjected to end repair and A-tailing using NEBNext Ultra II End Prep reagents (30 min at 20 °C followed by 30 min at 65 °C), followed by ligation of BAC-specific forked adapters for 30 min at 20 °C. The adapters were prepared by annealing complementary oligonucleotides at 98 °C followed by gradual cooling to 4 °C (1 °C per 15 s). Adapter and primer sequences are provided in the Supplementary Methods.

Adapter-ligated DNA was purified using AMPure XP beads at a 1:1 bead-to-sample volume ratio and quantified. A calibration PCR was performed using KAPA HiFi HotStart ReadyMix and 5′-biotinylated BAC-specific primers (see Supplementary Methods). Aliquots were collected after 6, 8, 10 and 12 PCR cycles and analyzed by 1% agarose gel electrophoresis to determine the optimal number of amplification cycles (1-2 cycles before reaching gel smear saturation). BAC-derived capture probes were then produced by three 50 µl PCR reactions using the conditions established during calibration. PCR products were pooled, purified using AMPure XP beads at a 1.7:1 bead-to-sample volume ratio, and quantified before use with DeNovix high-sensitivity assay.

### Micro-C chromatin fragmentation and library preparation

Frozen cell pellets containing 5 × 10⁶ cells were thawed on ice and lysed in ice-cold buffer containing 20 mM HEPES, 2 mM MgCl₂, 0.5× protease inhibitor cocktail, and 0.5× PMSF. Following 5 min on ice, 1% digitonin stock solution was added to a final concentration of 0.01%, followed by an additional 5 min on ice. Cell membranes were disrupted by repeated pipetting (10×) against the bottom of the microcentrifuge tube using a 1 mL pipette tip. Nuclei were centrifuged at 1,000 × g for 4 min at 4 °C and resuspended in MNase digestion buffer (10 mM Tris-HCl, pH 7.4, 1 mM CaCl₂). Chromatin was digested with micrococcal nuclease (MNase) for 1 h at 37 °C with agitation at 1,000 rpm. The MNase amount was calibrated on the wildtype sample to obtain predominantly mononucleosomal fragments, as assessed by 1% agarose gel electrophoresis. Calibration reactions tested 5–35 U of MNase per 110 µl reaction (see Supplementary Methods). Based on these results, we chose 70U for 450 µl reaction. Digestion was terminated by addition of EGTA to a final concentration of 5 mM.

Following MNase digestion, nuclei were washed with PBS and resuspended in ligation buffer containing dNTPs and ATP. DNA ends were repaired using 100 U of T4 polynucleotide kinase and 50 U of Klenow fragment, added simultaneously and incubated for 1 h at 37 °C with agitation at 1,000 rpm. Proximity ligation was initiated by adding 5,000 U of T4 DNA ligase and incubating for 1 h at 37 °C with agitation at 1,000 rpm, followed by addition of another 5,000 U of ligase and overnight incubation at 20 °C. Crosslinks were reversed by incubation with proteinase K at 65 °C for 2–3 h. DNA was purified by phenol–chloroform extraction and ethanol precipitation, followed by RNase A treatment.

To reduce the proportion of unligated mononucleosomal fragments, DNA was subjected to size selection using AMPure XP beads at a 0.7:1 bead-to-sample volume ratio, retaining the bead-bound DNA. DNA concentration was measured using a high-sensitivity DeNovix assay.

Sequencing libraries were prepared from 500 ng of size-selected DNA using the NEBNext Ultra II DNA Library Prep Kit. End repair and A-tailing were performed for 30 min at 20 °C followed by 30 min at 65 °C. Illumina adapters were ligated for 30 min at 20 °C, followed by USER enzyme treatment for 15 min at 37 °C. DNA was purified using AMPure XP beads at a 0.9:1 bead-to-sample volume ratio and resuspended in 27 µl 10 mM Tris-HCl. Libraries were amplified using KAPA HiFi HotStart ReadyMix and indexed primers from the NEBNext Multiplex Oligos for Illumina kit. The number of PCR cycles was determined by 25 µl calibration PCR with 5 µL of adapter ligated library as template, testing 4, 6, 8 and 10 cycles and selecting the minimal number of cycles producing detectable DNA product smear on a 1% agarose gel. Libraries were amplified in two 50 µl reactions per sample, pooled, purified using AMPure XP beads at a 1.7:1 bead-to-sample volume ratio, and quantified using a high-sensitivity DeNovix assay. Libraries from individual samples were subsequently pooled to a total amount of 1 µg DNA per one capture reaction.

### Targeted Enrichment of Micro-C Libraries Using BAC-Derived Probes

Targeted enrichment was performed using two successive rounds of hybridization with biotinylated BAC-derived probes. For the first round, a library mixture containing up to 1 µg of pooled Micro-C libraries, 3 µg of sheared salmon sperm DNA, and blocking oligonucleotides complementary to the library adapter sequences (20 pmol each) was prepared. Separately, a probe mixture containing 500 ng of biotinylated BAC-derived probes, 1.5 µg of Cot-1 DNA to block repetitive sequences, and blocking oligonucleotides complementary to the probe adapter sequences (10 pmol each) was prepared. Oligonucleotide sequences are provided in the Supplementary Methods.

The two mixtures were denatured separately at 95 °C for 5 min, followed by incubation at 65 °C for 10 min. Pre-warmed hybridization buffer was added to each mixture, followed by incubation at 65 °C for 1 h. The library and probe mixtures were then combined and incubated at 65 °C for 20–24 h. The final hybridization conditions were 5× SSPE, 5× Denhardt’s solution, 5 mM EDTA and 0.1% SDS.

Following hybridization, the samples were incubated for 30 min at room temperature with 15 µl of streptavidin-coated magnetic beads, pre-washed in high salt buffer (10 mM Tris-HCl, pH 7.5, 1 M NaCl, 1 mM EDTA), with occasional mixing. The beads with pooled down library fragments were washed once with low stringency wash buffer (1× SSC, 0.1% SDS) for 15 min at room temperature, then three times with high stringency wash buffer (0.1× SSC, 0.1% SDS) for 10 min each at 65 °C with shaking at 800 rpm, and three times with low salt wash buffer (10 mM Tris-HCl, pH 7.5, 10 mM NaCl). After the last wash, beads were resuspended in 26 µl of 10 mM Tris-HCl.

The captured libraries were amplified directly from the beads (5 µl of bead suspension per reaction) using KAPA HiFi HotStart ReadyMix and Illumina PCR primers in four 50 µl reactions. To determine the optimal number of PCR cycles, one calibration PCR reaction of 25 µl was performed with aliquots of PCR products taken after 16, 18, 20, and 22 cycles. The final amplification was then carried out using 1-2 cycles fewer than the number required for the DNA product smear on the gel to reach saturation. Amplified products were pooled, purified using AMPure XP beads at a 1.7:1 bead-to-sample volume ratio and quantified using a high-sensitivity DeNovix assay.

The amplified library was subjected to a second round of hybridization using 1 µg of the first-capture library, fresh BAC-derived probes (500 ng) and the same hybridization and washing conditions; after the final wash beads were resuspended in 26 µl of 10 mM Tris-HCl. Following the second capture, the optimal number of PCR cycles was determined by 25 µl calibration PCR with 5 µl of bead suspension testing 8, 10, 12 and 14 cycles; final amplification was then carried out in 2 50 µl PCR reactions using 1-2 cycles fewer than the number required for the smear on the gel to reach saturation. Amplified products were pooled, purified using AMPure XP beads at a 1.7:1 bead-to-sample volume ratio, and quantified using a high-sensitivity DeNovix assay.

The final libraries were sequenced 2×150 bp by paired-end sequencing.

### C-TALE

The C-TALE experiment was conducted as previously described^30^, but following Hi-C^45^ rather than 3C-Seq. Briefly, HAP1 cells were crosslinked, permeabilized, and digested with DpnII. In contrast to the published C-TALE protocol, which does not include biotin fill-in and streptavidin pull-down during this stage, DNA ends were filled in with biotin-14-dATP using DNA polymerase I Klenow fragment, followed by blunt-end ligation with T4 DNA ligase. Crosslinking was reversed by incubation with Proteinase K and heating, and DNA was purified by phenol–chloroform extraction and ethanol precipitation. Biotin was removed from unligated DNA ends using T4 DNA polymerase. DNA was then sonicated to a fragment size of 200–300 bp using a Covaris sonicator. Biotinylated ligation products were enriched using streptavidin-coated magnetic beads. The enriched DNA was subjected to end repair and A-tailing using the NEBNext Ultra II DNA Library Prep Kit, followed by Illumina adaptors ligation. Libraries were amplified by PCR using indexed primers, purified with AMPure XP beads, and quantified. Subsequent BAC-based probe preparation and target enrichment were performed as described for Micro-C libraries.

### Generation of CTCF insertion mutant

To introduce a CTCF binding site into the targeted genomic region, HAP1 cells were edited using CRISPR-Cas9. Cells were cultured in Iscove’s Modified Dulbecco’s Medium (IMDM) supplemented with 10% fetal bovine serum and 1% penicillin–streptomycin, and transfected at approximately 60% confluence.

Cells were co-transfected with 2.5 µg of a pSpCas9(BB)-2A-GFP (PX458) plasmid expressing Cas9 and guide RNA 313, targeting the insertion site (Chr1:30,817,713), and 2.5 µg of a double-stranded DNA repair template containing the CTCF binding site flanked by 200 bp homology arms. The inserted sequence was taken from Chr2:43,056,093-43,056,234. The repair template was excised from a custom pUC57-based plasmid (GenScript) using EcoRI and HindIII, and purified before transfection. Transfection was performed using jetOPTIMUS reagent (Polyplus) according to the manufacturer’s instructions. Guide RNA and PCR primer sequences are provided in the Supplementary Methods. At 24 h after transfection, cells were harvested, passed through a 40 µm strainer, and subjected to fluorescence-activated cell sorting. Individual GFP-positive cells were sorted into 96-well plates containing 200 µl IMDM per well, and clonal outgrowth was monitored for two weeks.

Genomic DNA was extracted from expanded clones using either alkaline lysis or the Norgen Cells and Tissue DNA Isolation Kit. Clones were screened by PCR using primers flanking the insertion site (CRM 1A F/R or CRM 2C F/R). The expected amplicon sizes were 624 bp for the knock-in allele and 409 bp for the wildtype allele. One clone yielded both the wildtype and knock-in PCR products, consistent with the presence of both alleles (HAP1 cells often go through diploidization^44^). Sanger sequencing of the knock-in PCR product using primer CRM 1A F confirmed integration of the donor sequence at the targeted locus, without additional sequence alterations in the sequenced amplicon.

### Data analysis

Micro-C-TALE, C-TALE and Micro-C data libraries were processed using distiller-nf (https://github.com/open2c/distiller-nf) and cooler^46^ to generate interaction maps in mcool format. Briefly, paired-end reads were aligned to the human genome (hg38), uniquely mapping non-duplicate read pairs were retained, reads were binned into specified resolutions and interaction maps were filtered and balanced. For Micro-C-TALE and C-TALE, balancing was performed only on the submatrix containing the target region. Micro-C data was taken from Barshad et al.^37^ and Hi-C data was taken from Guetta et al.^45^. CTCF ChIP-Seq was taken from Haarhuis et al.^43^ using Cistrome Data Browser^47^. On target read pairs were defined as read pairs in which both sides map to the target region.

## Data availability

Raw and processed Micro-C-TALE and C-TALE data files have been submitted to GEO and are available at GSEXXXXX.

## Acknowledgements

We thank the Azrieli Technion Genomic Center, David Cohen, as well as past and present members of the Kaplan lab. This research was supported by Israel Science Foundation Personal Grant (3599/25) (NK), Technion funding (AKG) and a scholarship by The Iris and Winifred Wolfson Endowment Fund via the Administrator General (AG).

