## Supplementary Methods for "Micro-C-TALE: BAC-based targeted high-resolution measurement of 3D chromatin interactions"

A. Micro-C
B. MNase calibration
C. Preparation of forked adapters for probe amplification
D. BAC probes production
E. Capture
F. Oligo and primer sequences
G. Reagents and equipment

### A. Micro-C

#### 1. Cells grow and fixation:

1.1. Grow cells in a big flask until confluency.
1.2. Harvest cells with trypsin and count.
1.3. Centrifuge 5 min, 500g, RT.
1.4. Resuspend cells with 950 µl PBSX1 for every 10M cells.
1.5. Add 55 µl of 37% Formaldehyde for every 10M cells.
1.6. Incubate 10 mins on rocking platform.
1.7. Add dropwise 600 µl 2M Tris-HCl pH 7.4 for every 10M cells.
1.8. Rock for 5 mins.
1.9. Put on ice for 10 mins.
1.10. Centrifuge at 1000g, +4 C°, for 5 min.
1.11. Remove supernatant.
1.12. Resuspend the cells with 950 µl PBSX1 for every 10M cells.
1.13. Add 50 µl 2M Tris-HCl for every 10M cells.
1.14. Split to aliquots of 5M cells each.
1.15. Centrifuge at 1000g, +4C° for 5 min.
1.16. Remove supernatant.
1.17. Snap-freeze the cell pellets in liquid nitrogen, store at -80C°.

#### 2. Before the experiment

2.1. Turn on +4C° small centrifuge.
2.2. Turn on +37C° thermoshaker.
2.3. Bring ice.
2.4. Thaw PIC and PMSF, vortex.
2.5. Thaw Digitonin 1%.
2.6. Prepare Stock Lysis buffer:

| | **Stock** | **Cf** | **V** |
|---|---|---|---|
| Hepes | 1M | 20 mM | 200 µl |
| MgCl2 | 1M | 2mM | 20 µl |
| PMSF 100mM | 100X | 0.5X | - |
| PIC 100mM | 100X | 0.5X | - |
| UPW | | | 9.78 ml |
| Total | | | 10 ml |

(PMSF and PIC: Add fresh before the experiment)

2.7. Prepare MNase Buffer:

| | **Stock** | **Cf** | **V** |
|---|---|---|---|
| Tris-HCl pH 7.4 | 2M | 10 mM | 50 µl |
| CaCl2 | 1M | 1mM | 10 µl |
| UPW | | | 9.95 ml |
| Total | | | 10 ml |

#### 3. Lysis

3.1. Prepare Lysis reaction mix in low binding tube, place on ice.

| | **V** | |
|---|---|---|
| Stock Lysis Buffer | 500 µl | |
| PIC | 2.5 µl | |
| PMSF | 2.5 µl | |
| | | 505µl/tube |

3.2. Thaw the frozen cells on ice for a few minutes.

3.3. Resuspend cells with ~350 µl Lysis mix (using a low-binding tip) and transfer them back to the lysis reaction tube (keep the tip).

3.4. Incubate 5 mins on ice.

3.5. Add 5 µl of 1% Digitonin, mix with the same low-binding tip, incubate 5 mins on ice.

3.6. Do 10 ups-and-downs of all the volume against the tube walls (using the same tip).

3.7. Centrifuge for 4 mins in +4C° at 1000 g.

3.8. Discard the liquid carefully.

#### 4. MNase digestion

4.1. Add to the pellet 500 µl MNase buffer, vortex.

4.2. Take out 50 µl aliquot for Chromatin Integrity control.

4.3. To the rest 450 µl add ________ of MNase, vortex.

*The right amount of MNase should be calibrated for each fixation batch, using MNase calibration protocol.

4.4. Incubate on thermoshaker for 1h at 37C°, 1000rpm.

4.5. To CI control add and mix:

| | V | |
|---|---|---|
| PBSX1 | 175 µl | |
| EGTA 0.5M | 1 µl | |
| SDS 10% | 25 µl | |
| | | 201µl/tube |

#### 5. Ligation

5.1. Thaw: 10mM dNTPs, 10mM ATP, T4 DNA Ligase buffer.
5.2. After 1h add 4.5 µl of EGTA 0.5M, vortex.
5.3. Centrifuge for 5 mins, 500g, RT.
5.4. Discard liquid, add 500 µl PBSX1, vortex.
5.5. Centrifuge for 5 mins, 1000g, RT.
5.6. Prepare Ligation reaction mix and distribute to new eppendorfs:

| | **Stock** | **Cf** | **V X1** | |
|---|---|---|---|---|
| T4 DNA Ligase Buffer | 10X | 1.05X | 48.3 µl | |
| dNTPs | 10 mM | 0.105 mM | 4.83 µl | |
| ATP | 10 mM | 2.1 mM | 96.6 µl | |
| UPW | | | 310 µl | |
| total | | | 460 µl | 460 µl/tube |

5.7. Discard the liquid.
5.8. Resuspend the cells with ~350 µl Ligation mix and transfer them back to the ligation reaction tube.
5.9. Take out 50 µl aliquot for Digestion Control. Add and mix:

| | V | |
|---|---|---|
| PBSX1 | 175 µl | |
| SDS 10% | 25 µl | |
| | | 200 µl/tube |

5.10. To the rest 410 µl add: 100U T4 PNK = 10 µl; 50U Klenow = 10 µl. Vortex.
5.11. Incubate on thermoshaker for 1h, at 37C°, 1000 rpm.
5.12. After 1h add 5000U T4 DNA ligase = 12.5 µl, vortex.
5.13. Incubate on thermoshaker for 1h, at 37C°, 1000 rpm.
5.14. Add again 5000U T4 DNA ligase = 12.5 µl, vortex.

5.15. Incubate overnight on thermoshaker at 20 C°, 1000 rpm.
5.16. To CI and D control tubes add 2 µl Proteinase K 800U/ml, incubate on thermoshaker at 65C° overnight.
*Seal with parafilm or use o-ring tubes.
5.17. Divide the ligation reaction to 2 tubes of 225 µl.
5.18. Add 25 µl of SDS10% to each ligation tube.
5.19. Add 3 µl Proteinase K 800U/ml to each ligation tube.
5.20. Incubate on thermoshaker at 65C° for 2-3h.

#### 6. DNA extraction (CI&D controls and LigationX2)

6.1. Turn on +4C° centrifuge.
6.2. Take out Phenol-Chloroform to warm to RT.
6.3. Add 1V (250 µl) of Phe-Cl to each tube, vortex.
6.4. Centrifuge at 17,000g, RT, 3.5 min.
6.5. Move the upper liquid phase to a new tube (210 µl).
6.6. Add 1/15V (15 µl) Sodium Acetate 3M pH 5.2 to each tube.
6.7. Add 3V (750 µl) of cold 96% EtOH, vortex.
6.8. Centrifuge at 17,000g, +4C°, 10 min.
6.9. Discard the liquid to a 2ml tube to avoid losing the pellet.
6.10. Add 400 µl RT 80% EtOH.
6.11. Centrifuge at 17,000g, +4C°, 5 min.
6.12. Discard the liquid to a 2ml tube, dry with vacuum.
6.13. Mix:

| | X1 |
|---|---|
| Tris-HCl 10 mM | 150 µl |
| RNaseA | 4 µl |

6.14. Add 25 µl to control tubes. Add 100 µl to first ligation tube and transfer it to the second ligation tube.
6.15. Incubate for 10 min at 37C°.
6.16. Measure concentration with HS Denovix.
It is important to correctly measure Digestion control and Ligation, to load same amount of DNA to the gel.
6.17. Run ~200ng of DNA on 1% Agarose gel.
The mono-nucleosome band should be less visible in ligation in favor of higher bands.

7. Enrichment of ligated products

7.1. Add Tris-HCl 10 mM to ligation tube to bring volume back to 100 µl.
7.2. Warm Ampure XP the beads 30 mins before use and resuspend thoroughly.
7.3. Add 0.7V (70 µl) to the tube and mix well.
7.4. Incubate 10 mins at RT, place on MPS, wait until beads stick to the magnet.
7.5. Transfer the liquid (mono-nucleosome - MN) to a new tube or discard.
7.6. Wash the beads twice with 80% EtOH. After 2$^{nd}$ time incubate 5 mins. Air dry the beads.
7.7. Resuspend in 30 µl Tris-HCl 10mM.
7.8. Incubate 5 mins at RT, put on MPS, wait until beads stick to the magnet.
7.9. Transfer to a new tube (Ligation -L).
7.10. Measure concentration with HS Denovix.
7.11. Run 150 ng of DNA on 1% Agarose gel.
7.12. Continue to library preparation with 500 ng using NEBNext kit.

8. NEBNext End Prep

8.1. Start PCR program before mixing to cool down the lid to 75C°.
8.2. Mix on ice:

| | X1 | X |
|---|---|---|
| NEBNext Ultra II End Prep Enzyme Mix | 3 µl | |
| NEBNext Ultra II End Prep Reaction Buffer | 7 µl | |
| ligated DNA+UPW | 50 µl | - |
| Total | 60 µl | 10 µl/tube |

*check for sediment and dissolve it

8.3. Mix well the entire volume by pipetting.
8.4. The program is 20°C for 30 min to add adenine on the 3' end and then 65°C for 30 min to inactivate enzymes with heated lid set to 75°C.
8.5. Place tubes on ice and proceed to adaptor ligation immediately.

#### 9. Adaptor ligation

9.1. Start PCR program on a different head before mixing to cool down the lid to 47°C.
9.2. Mix on ice:

| | X1 | X |
|---|---|---|
| End Prep Reaction Mixture | 60 µl | |
| (red) NEBNext Adaptor for Illumina | 2.5 µl | |
| (red) NEBNext Ultra II Ligation Master Mix | 30 µl | |
| (red) NEBNext Ligation Enhancer | 1 µl | |
| | 93.5 µl | |

9.3. Mix well the entire volume by pipetting.
9.4. The program is 20°C for 30 min with heated lid set to 47°C.
9.5. Add 3 µl of (red or blue) USER® Enzyme to the ligation mixture.
9.6. Mix well and incubate at 37°C for 15 minutes with the heated lid set to 47°C.

#### 10. Cleanup with Ampure XP beads

10.1. Warm the beads 30 mins before use and resuspend thoroughly.
10.2. Add 0.9 volumes (87 µl) to the tube, mix well.
10.3. Incubate for 10 min at RT, place on MPS, wait until beads stick to the magnet.
10.4. Discard the liquid.
10.5. Wash twice with 80% ethanol: I) up-and-downs 3-4 times, II) incubate for 5 min on MPS.
10.6. Discard the liquid, air-dry the beads for 5 mins.
10.7. Resuspend the beads in 27 µl Tris-HCl 10mM, mix well.
10.8. Incubate for 5 min at RT, put on MPS, wait until beads stick to the magnet.
10.9. Transfer the liquid to a new tube.
10.10. Check concentration with HS Denovix.

#### 11. Calibration PCR

11.1. Set up the reaction in UV cabinet:

*Write down the indexes used.

*Set up negative control as well, without DNA fragments

| | X1 |
|---|---|
| Kappa HiFi HS ready mix | 12.5 µl |
| Adaptor ligated DNA fragments | 5 µl |
| Index primers | 2.5 µl |
| UPW | 5 µl |
| total | 25 µl |

11.2. Run PCR program:

| Step | Temp | Time | |
|---|---|---|---|
| 1 | 95 C° | 3 min | |
| 2 | 98 C° | 20 sec | |
| 3 | 65 C° | 30 sec | X10 cycles |
| 5 | 72 C° | 45 sec | |
| 6 | 72 C° | 3 min | |
| 7 | 4°C | ∞ | |

11.3. Take 3 µl aliquots after 4,6, 8 and 10 cycles at the last 15 seconds of elongation.

(Negative control - only at the end)

11.4. Prepare gel loading mix: UPW 7 µl + Loading dye 2 µl, distribute 9 µl to each tube.

11.5. Run on 1% agarose gel. Choose the minimal cycle for amplification.

If there is a problem with the gel check concentration with HS Denovix.

11.6. Do not proceed to production PCR in the same day.

#### 12. Production PCR

12.1. Set up 2X50 µl reactions in UV cabinet:

*Write down the indexes used.

| | X1 | X2 |
|---|---|---|
| Kappa HiFi HS ready mix | 25 µl | 50 µl |
| Adaptor ligated DNA fragments | 10 µl | 20 µl |
| Index primers | 5 µl | 10 µl |
| UPW | 10 µl | 20 µl |
| total | 50 µl | 100 µl |

12.2. Run PCR program according to calibration:

| Step | Temp | Time |
|---|---|---|
| 1 | 95 C° | 3 min |
| 2 | 98 C° | 20 sec |
| 3 | 65 C° | 30 sec |
| 5 | 72 C° | 45 sec |
| 6 | 72 C° | 3 min |
| 7 | 4°C | ∞ |

12.3. Purify the reaction with Ampure XP beads:

- 12.3.1. Mix the two reactions in one eppendorf.
- 12.3.2. Add 1.7 volumes of Ampure beads (resuspend thoroughly) to the tube (170 µl).
- 12.3.3. Incubate for 10 min at RT. Place on MPS.
- 12.3.4. Wash twice with 80% ethanol: I) up-and-downs 3-4 times, II) incubate for 5 min on MPS.
- 12.3.5. Air-dry the beads.
- 12.3.6. Resuspend the beads in 20 µl Tris-HCl 10mM, incubate for 5 min, put on MPS.
- 12.3.7. Transfer the liquid to a new tube.
- 12.3.8. Check concentration with HS Denovix.

12.4. Prepare a mix of 1 µg from all the libraries and continue to capture.

### B. MNase calibration

1. Before the experiment
   1.1. Turn on +4C° small centrifuge
   1.2. Turn on +37C° thermoshaker
   1.3. Bring ice
   1.4. Thaw PIC and PMSF, vortex
   1.5. Prepare Stock Lysis buffer:

| | Stock conc. | Cf | V |
|---|---|---|---|
| Hepes | 1M | 20 mM | 200 µl |
| MgCl2 | 1M | 2mM | 20 µl |
| PMSF 100mM | 100X | 0.5X | - |
| PIC 100mM | 100X | 0.5X | - |
| UPW | | | 9.78 ml |
| Total | | | 10 ml |

} Add fresh before experiment

   1.6. Prepare MNase Buffer:

| | Stock conc. | Cf | V |
|---|---|---|---|
| Tris-HCl pH 7.4 | 2M | 10 mM | 50 µl |
| CaCl2 | 1M | 1mM | 10 µl |
| UPW | | | 9.95 ml |
| Total | | | 10 ml |

2. Prepare Lysis reaction: 500 µl Stock Lysis buffer + 2.5 µl PMSF + 2.5 µl PIC, place on ice.
3. Thaw 5M frozen cells on ice for a few minutes.
4. Resuspend cells with ~350 µl Lysis solution (using a low-binding tip) and transfer them back to the lysis reaction tube (keep the tip).
5. Incubate 5 mins on ice.
6. Add 5 µl of 1% Digitonin, mix with the same low-binding tip, incubate 5 mins on ice.
7. Do 10 ups-and-downs of all the volume against the tube walls (using the same tip).
8. Centrifuge 4 mins in +4C° at 1000 g.
9. Discard the liquid carefully.
10. Add to the precipitate 500 µl MNase buffer, vortex.
11. Take out 50 µl aliquot for Chromatin Integrity control.

12. Split the rest into 4 tubes of 110 µl each.
13. Digest with different MNase concentrations: *dilute the 20U/µl MNase to 5U/µl using MNase buffer.

| Units | 5 | 15 | 25 | 35 |
|---|---|---|---|---|
| Volume | 1 µl | 3 µl | 1.25 µl | 1.75 µl |

14. Vortex, incubate on thermoshaker for 1h at 37C°, 1000rpm.
15. To CI control add: 175 µl PBSX1, 1 µl EGTA 0.5M, mix; 25 µl SDS 10%.
16. After 1h add 1.1 µl of EGTA 0.5M, vortex.
17. Add 115 µl PBSX1 and 25 µl SDS 10%, vortex.
18. To all digestion and CI tubes add ProteinaseK 800U/ml:
    2 µl to control; 3 µl to digestion
19. Incubate at 65C° for 2-3h,1000 rpm on thermoshaker (can be done overnight if sealed with parafilm).
20. Take out Phe-Cl from +4C° to warm to RT.
21. Add 1V (250 µl) of Phe-Cl to each tube, vortex.
22. Centrifuge at 17,000g, +4C°, 3.5 min.
23. Move the upper liquid phase to a new tube (210 µl).
24. Add 1/15V (15 µl) Sodium Acetate 3M pH 5.2 to each tube.
25. Add 3V (750 µl) of cold 96% EtOH, vortex.
26. Centrifuge at 17,000g, +4C°, 10 min.
27. Discard the liquid to a 2ml tube to avoid losing the precipitate.
28. Add 400 µl RT 80% EtOH.
29. Centrifuge at 17,000g, +4C°, 5 min.
30. Discard the liquid to a 2ml tube, dry with vacuum.
31. Mix: 150 µl Tris-HCl 10mM + 4 µl RNaseA.
32. Add 25 µl to each tube, incubate for 10 min at 37C°.
33. Measure concentration with HS Denovix.
34. Run ~200ng on 1% Agarose gel.

Look for a mostly mono-nucleosome band but without a smear under it. Some bands of higher m.w. should be visible as well.

#### C. Preparation of forked adapters for probe amplification

1. Prepare BAC-adapter annealing reaction in a PCR tube as follows:

| Ingredient | Volume |
|---|---|
| 10× NEBuffer 2.1 | 10 µl |
| BAC-adapter-for (100 µM) | 10 µl |
| BAC-adapter-rev (100 µM) | 10 µl |
| milliQ | 70 µl |

2. Place the tube in a thermal cycler, heat the mix to 98 °C, then gradually cool to 4 °C at a rate of 1 °C per 15 s. Do not warm the tube to RT.
3. Dilute annealed Y-adapters 10 times with cold 10 mM Tris-HCl (pH 8.0), split in 10 aliquots, and store at −20 °C.

#### D. BAC probes production

1. DNA Sonication

    1.1. Take 1300 ng BAC DNA in 130 µl UPW and transfer to a Covaris microTUBE for sonication.
    *It is possible to use several BACs or other plasmids together mixed in an equimolar quantity.
    1.2. Shear the DNA to a size of 200-300 bp using a sonicator.

2. NEB End-Repair

    2.1. Thaw Reaction buffer.
    2.2. Bring ice.
    2.3. Start PCR program before mixing to cool down the lid to 75C°.
    2.4. Mix End-Repair components in PCR tube on ice:

| Ingredient | X1 | |
|---|---|---|
| Sonicated BAC DNA | 50 µl | |
| NEBNext Ultra II End Prep Reaction buffer | 7 µl | *check for sediment and dissolve it |
| NEBNext Ultra II End Prep Enzyme Mix | 3 µl | |
| TOTAL | 60 µL | |

    2.5. Mix well the entire volume by pipetting.
    2.6. The program is 20°C for 30 min to add adenine on the 3' end and then 65°C for 30 min to inactivate enzymes with heated lid set to 75°C.
    2.7. Place tubes on ice and proceed to adaptor ligation immediately.

3. Adaptors Ligation

    3.1. Start_PCR program on a different head before mixing to cool down the lid to 47°C.
    3.2. Prepare Adaptors ligation mix:

| Ingredient | X1 |
|---|---|
| End Repair Reaction Mixture | 60 µl |
| BAC adaptors [5 µM] | 2.5 µl |
| NEBNext Ultra II Ligation Master Mix | 30 µl |
| NEBNext Ligation Enhancer | 1 µl |
| Total | 93.5 µl |

3.3. Mix well the entire volume by pipetting.

3.4. The program is 20°C for 30 min with heated lid set to 47°C.

4. Clean up with Ampure XP beads:

4.1. Warm the beads 30 mins before use and resuspend thoroughly.

4.2. Add X1 volumes (93.5 µl) to the tube, mix well.

4.3. Incubate for 10 min at RT, place on MPS, wait until beads stick to the magnet.

4.4. Discard the liquid.

4.5. Wash twice with 80% ethanol: I) up-and-downs 3-4 times, II) incubate for 5 min on MPS.

4.6. Discard the liquid, air-dry the beads for 5 mins.

4.7. Resuspend the beads in 50 µl Tris-HCl 10mM, mix well.

4.8. Incubate for 5 min at RT, put on MPS, wait until beads stick to the magnet.

4.9. Transfer the liquid to a new tube, mark properly.

4.10. Check concentration with HS Denovix.

4.11. Keep this tube in -80°C for any period of time.

5. Calibration PCR

5.1. Prepare PCR calibration reaction in the UV cabinet:

| Ingredient | X1 |
|---|---|
| Kappa HiFi HS ready mix | 12.5 µl |
| Adaptor Ligated BAC DNA | 2 µl |
| BACbioFOR Primer [10 µM] | 1.25 µl |
| BACbioREV Primer [10 µM] | 1.25 ul |
| UPW | 8 µl |
| TOTAL | 25 µL |

5.2. Run PCR program:

| Step | Temp | Time | |
|---|---|---|---|
| 1 | 95 C° | 3 min | |
| 2 | 98 C° | 20 sec | X12 |
| 3 | 65 C° | 30 sec | |
| 5 | 72 C° | 45 sec | |
| 6 | 72 C° | 3 min | |
| 7 | 4°C | ∞ | |

5.3. Take 3 µl aliquots after 6,8, 10 and 12 cycles at the last 15 seconds of elongation.
5.4. Prepare gel loading mix: UPW 7 µl + Loading dye 2 µl, distribute 9 µl to each tube.
5.5. Run on 1% agarose gel.
5.6. Check concentration of the end product with High Sensitivity Denovix.
5.7. Do not proceed to production PCR in the same day.

#### 6. Production PCR

6.1. Set up 3X50 µl production PCR reactions based on results from the previous section:
6.2. Mix in the UV cabinet:

| Ingredient | X1 | X3 |
|---|---|---|
| Kappa HiFi HS ready mix | 25 µl | 75 µl |
| Adaptor Ligated BAC DNA | 4 µl | 12 µl |
| BACbioFOR Primer [10 µM] | 2.5 µl | 7.5 µl |
| BACbioREV Primer [10 µM] | 2.5 µl | 7.5 µl |
| UPW | 16 µl | 48 µl |
| TOTAL | 50 µL | 150 µl |

6.3. Run PCR program:

| Step | Temp | Time | |
|---|---|---|---|
| 1 | 95 C° | 3 min | |
| 2 | 98 C° | 20 sec | X |
| 3 | 65 C° | 30 sec | |
| 5 | 72 C° | 45 sec | |
| 6 | 72 C° | 3 min | |
| 7 | 4°C | ∞ | |

7. Clean up with Ampure XP beads:

7.1. Warm the beads 30 mins before use and resuspend thoroughly.
7.2. Mix the 3 reactions to one low binding 1.5 ml eppendorf.
7.3. Add X1.7 volumes (255 µl) to the tube, mix well.
7.4. Incubate for 10 min at RT, place on MPS, wait until beads stick to the magnet.
7.5. Discard the liquid.
7.6. Wash twice with 80% ethanol: I) up-and-downs 3-4 times, II) incubate for 5 min on MPS.
7.7. Discard the liquid, air-dry the beads for 5 mins.
7.8. Resuspend the beads in 25 µl Tris-HCl 10mM, mix well.
7.9. Incubate for 5 min at RT, put on MPS, wait until beads stick to the magnet.
7.10. Transfer the liquid to a new tube, mark properly.
7.11. Check concentration with HS Denovix.

### E. Capture

1. Hybridization 1
   1.1. Prepare libraries mix:

| library | | | | | | total V |
|---|---|---|---|---|---|---|
| C [ng/ µl] | | | | | | |
| V for 1 reaction | | | | | | |
| V for ___ reactions | | | | | | |

   1.2. Prepare Hybridization buffer:

| | Stock | V | X |
|---|---|---|---|
| SSPE buffer | 20X | 250 µl | |
| Denhardt's solution | 50X | 100 µl | |
| EDTA | 0.5 M | 10 µl | |
| SDS | 10% | 10 µl | |
| UPW | | 130 µl | |
| total | | 500 µl | |

*Warm mixture to 65C° before addition of 20XSSPE to avoid precipitate formation.

   1.3. Place the thermoshaker with the hybridization mix in close proximity to PCR.
   1.4. Prepare two mixes in PCR tubes as follows:

*To increase the complexity, prepare several hybridization reactions. Once the mixes are ready, split them into individual PCR tubes.

*For more than 3 different BACs use 1 µg probes, 3 µl Cot, 2 µl primers.

1.4.1. Library mix

| | | X4.2 |
|---|---|---|
| Libraries | Up to 1 µg | |
| Sheared salmon DNA(1 µg/µl solution in UPW) | 3 µl | |
| Illumina Uni oligo (10 µM) | 2 µl | |
| Illumina rev(10 µM) | 2 µl | |
| UPW | Up to 30 µl | |

1.4.2. Probe mix

| | | X4.2 |
|---|---|---|
| Probes | 500 ng | |
| Cot-1 DNA (1 µg/µl) | 1.5 µl | |
| BAC-adapter-for (10 µM) | 1 µl | |
| BAC-rev primer (10 µM) | 1 µl | |
| UPW | Up to 30 µl | |

1.5. Place each tube in the PCR and incubate: 5  mins at 95 C° and 10 mins at 65 C°.
1.6. Add 30 µl hybridization mix to each tube, on the PCR to avoid cooling down.
1.7. Incubate 1h at 65 C°.
1.8. Transfer the Probe mix to the Library mix.
1.9. Incubate for 20-24h at 65 C°.

2. Buffers to prepare before pulldown:

2.1. WB buffer (store at +4C°)

| | Stock | Cf | V |
|---|---|---|---|
| Tris-HCl pH7.5 | 1 M | 10 mM | 10 µl |
| NaCl | 5 M | 1 M | 200 µl |
| EDTA | 0.5 M | 1 mM | 2 µl |
| UPW | | | 788 µl |
| total | | | 1 ml |

2.2. LS buffer (store at +4C°)

| | Stock | Cf | V |
|---|---|---|---|
| SSC | 2X | 1X | 500 µl |
| SDS | 10% | 0.1% | 10 µl |
| UPW | | | 490 µl |
| total | | | 1 ml |

2.3. HS buffer (store at +4C°)

| | Stock | Cf | V |
|---|---|---|---|
| SSC | 2X | 0.1X | 50 µl |
| SDS | 10% | 0.1% | 10 µl |
| UPW | | | 940 µl |
| total | | | 1 ml |

2.4. S buffer (store at +4C°)

| | Stock | Cf | V |
|---|---|---|---|
| Tris-HCl pH7.5 | 1 M | 10 mM | 10 µl |
| NaCl | 5 M | 10 mM | 2 µl |
| UPW | | | 988 µl |
| total | | | 1 ml |

3. Biotin pulldown 1

*Combine all the reaction tubes into one. Adjust the volumes proportionally according to the number of reactions, up until the LS buffer stage. At this stage, regardless of the number of reactions, increase the volume to 500 µl (instead of the 300 µl used for a single reaction).

3.1. Add 200 µl WB buffer to low binding eppedorf.
3.2. Add 15 µl Streptavidin beads, put on MPS.
3.3. Discard the liquid, add 200 µl WB buffer.
3.4. Discard the liquid, add 300 µl WB buffer, remove from MPS.
3.5. Take the hybridization tube from PCR, spin down, add it to the beads, vortex, spin down.
3.6. Incubate 30 mins at RT, shake once every 10 mins.
(Meanwhile heat the thermoshaker near UV cabinet to 65 C° and prewarm HS buffer.)
3.7. Place on MPS, discard the liquid, remove from MPS.
3.8. Add 300 µl LS buffer, vortex.
3.9. Incubate 15 min at RT, shake occasionally. Spin down.
3.10. Place on MPS, discard the liquid.

*From now on work in UV cabinet, clean the MPS with ethanol and switch gloves.

3.11. Resuspend the beads with 300 µl prewarmed HS buffer and transfer them to new low binding eppendorf.
3.12. Incubate on thermoshaker for 10 min at 65 C°, 800 rpm.
3.13. Place on MPS, discard the liquid.
3.14. Resuspend the beads with 300 µl prewarmed HS buffer.
3.15. Incubate on thermoshaker for 10 min at 65 C°, 800 rpm.
3.16. Place on MPS, discard the liquid.
3.17. Resuspend the beads with 300 µl prewarmed HS buffer.
3.18. Incubate on thermoshaker for 10 min at 65 C°, 800 rpm.
3.19. Place on MPS, discard the liquid.
3.20. Resuspend in 200 µl S buffer and transfer to a new low binding eppendorf. Put on MPS.
3.21. Discard liquid, wash with 200 µl S buffer (3-4 up-and-downs).
3.22. Discard the liquid, add 200 µl S buffer, incubate few minutes on MPS.
3.23. Discard the liquid.
3.24. Resuspend the beads in 26 µl Tris-HCl 10 mM, transfer to PCR tube.

4. Calibration PCR 1

4.1. Set up the reaction in UV cabinet:

*Set up negative control as well, without DNA fragments

| | X1 |
|---|---|
| DNA library | 2.5 µl |
| Kappa HiFi HS ready mix | 12.5 µl |
| Ilumina For | 1.25 µl |
| Ilumina Rev | 1.25 µl |
| UPW | 7.5 µl |
| total | 25 µl |

4.2. Run PCR program:

| Step | Temp | Time | |
|---|---|---|---|
| 1 | 95 C° | 3 min | |
| 2 | 98 C° | 20 sec | X22 cycles |
| 3 | 65 C° | 30 sec | |
| 5 | 72 C° | 45 sec | |
| 6 | 72 C° | 3 min | |
| 7 | 4°C | ∞ | |

4.3. Take 3 µl aliquots after 16,18, 20 and 22 cycles at the last 15 seconds of elongation.

(Negative control - only at the end)

4.4. Prepare gel loading mix: UPW 7 µl + Loading dye 2 µl, distribute 9 µl to each tube.

4.5. Run on 1% agarose gel. Choose the minimal cycle for amplification.

If there is a problem with the gel check concentration with HS Denovix.

4.6. Do not proceed to production PCR in the same day.

5. Production PCR 1

5.1. Set up in UV cabinet 4X50 µl reactions:

| | X1 | X4 |
|---|---|---|
| DNA library | 5 µl | 20 µl |
| Kappa HiFi HS ready mix | 25 µl | 100 µl |
| Ilumina For | 2.5 µl | 10 µl |
| Ilumina Rev | 2.5 µl | 10 µl |
| UPW | 15 µl | 60 µl |
| Total | 50 µl | 200 µl |

5.2. Run PCR program according to calibration.

| Step | Temp | Time |
|---|---|---|
| 1 | 95 C° | 3 min |
| 2 | 98 C° | 20 sec |
| 3 | 65 C° | 30 sec |
| 5 | 72 C° | 45 sec |
| 6 | 72 C° | 3 min |
| 7 | 4°C | ∞ |

5.3. Purify the reaction with Ampure XP beads:

- 5.3.1. Combine all reactions to the same tube.
- 5.3.2. Add 1.7 volumes of Ampure beads (resuspend thoroughly) to the tube (340 µl).
- 5.3.3. Incubate for 10 min at RT. Place on MPS.
- 5.3.4. Wash twice with 80% ethanol: I) up-and-downs 3-4 times, II) incubate for 5 min on MPS.
- 5.3.5. Air-dry the beads.
- 5.3.6. Resuspend the beads in 25 µl Tris-HCl 10mM, incubate for 5 min, put on MPS.
- 5.3.7. Transfer the liquid to a new tube.
- 5.3.8. Check concentration with HS Denovix.

6. Hybridization 2

6.1. Prepare Hybridization buffer:

| | Stock | V |
|---|---|---|
| SSPE buffer | 20X | 500 µl |
| Denhardt's solution | 50X | 200 µl |
| EDTA | 0.5 M | 20 µl |
| SDS | 10% | 20 µl |
| UPW | | 260 µl |
| total | | 1000 µl |

*Warm mixture to 65C° before addition of 20XSSPE to avoid precipitate formation.

6.2. Place the thermoshaker with the hybridization mix in close proximity to PCR.

6.3. Prepare two mixes in PCR tubes as follows:

*To increase the complexity, prepare several hybridization reactions. Use all the capture 1 library. Once the mixes are ready, split them into individual PCR tubes.

6.3.1. Library mix

| | | X |
|---|---|---|
| Capture1 library | 1 µg | |
| Sheared salmon DNA(1 µg/µl solution in UPW) | 3 µl | |
| Illumina Uni oligo (10 µM) | 2 µl | |
| Illumina rev (10 µM) | 2 µl | |
| UPW | Up to 30 µl | |

6.3.2. Probe mix

| | | X |
|---|---|---|
| Probes | 1 µg | |
| Cot-1 DNA (1 µg/µl) | 3 µl | |
| BAC-adapter-for (10 µM) | 2 µl | |
| BAC-rev primer (10 µM) | 2 µl | |
| UPW | Up to 30 µl | |

1.1. Place each tube in the PCR and incubate: 5 mins at 95 C° and 10 mins at 65 C°.

1.2. Add 30 µl hybridization mix to each tube, on the PCR to avoid cooling down.

1.3. Incubate 1h at 65 C°.

1.4. Transfer the Probe mix to the Library mix.

1.5. Incubate for 20-24h at 65 C°.

2. Biotin pulldown
   *Combine all the reaction tubes into one. Adjust the volumes proportionally according to the number of reactions, up until the LS buffer stage. At this stage, regardless of the number of reactions, increase the volume to 500 µl (instead of the 300 µl used for a single reaction).

   2.1. Add 200 µl WB buffer to low binding eppedorf.
   2.2. Add 15 µl Streptavidin beads, put on MPS.
   2.3. Discard the liquid, add 200 µl WB buffer.
   2.4. Discard the liquid, add 300 µl WB buffer, remove from MPS.
   2.5. Take the hybridization tube from PCR, spin down, add it to the beads, vortex, spin down.
   2.6. Incubate 30 mins at RT, shake once every 10 mins.
   (Meanwhile heat the thermoshaker near UV cabinet to 65 C° and prewarm HS buffer.)
   2.7. Place on MPS, discard the liquid, remove from MPS.
   2.8. Add 300 µl LS buffer, vortex.
   2.9. Incubate 15 min at RT, shake occasionally. Spin down.
   2.10. Place on MPS, discard the liquid.

   *From now on work in UV cabinet, clean the MPS with ethanol and switch gloves.
   2.11. Resuspend the beads with 300 µl prewarmed HS buffer and transfer them to new low binding eppendorf.
   2.12. Incubate on thermoshaker for 10 min at 65 C°, 800 rpm.
   2.13. Place on MPS, discard the liquid.
   2.14. Resuspend the beads with 300 µl prewarmed HS buffer.
   2.15. Incubate on thermoshaker for 10 min at 65 C°, 800 rpm.
   2.16. Place on MPS, discard the liquid.
   2.17. Resuspend the beads with 300 µl prewarmed HS buffer.
   2.18. Incubate on thermoshaker for 10 min at 65 C°, 800 rpm.
   2.19. Place on MPS, discard the liquid.
   2.20. Resuspend in 200 µl S buffer and transfer to a new low binding eppendorf. Put on MPS.
   2.21. Discard liquid, wash with 200 µl S buffer (3-4 up-and-downs).
   2.22. Discard the liquid, add 200 µl S buffer, incubate few minutes on MPS.
   2.23. Discard the liquid.
   2.24. Resuspend the beads in 26 µl Tris-HCl 10 mM, transfer to PCR tube.

3. Calibration PCR 2

3.1. Set up the reaction in UV cabinet:

*Set up negative control as well, without DNA fragments

| | X1 |
|---|---|
| DNA library | 5 µl |
| Kappa HiFi HS ready mix | 12.5 µl |
| Ilumina For | 1.25 µl |
| Ilumina Rev | 1.25 µl |
| UPW | 5 µl |
| total | 25 µl |

3.2. Run PCR program:

| Step | Temp | Time | |
|---|---|---|---|
| 1 | 95 C° | 3 min | |
| 2 | 98 C° | 20 sec | X14 cycles |
| 3 | 65 C° | 30 sec | |
| 5 | 72 C° | 45 sec | |
| 6 | 72 C° | 3 min | |
| 7 | 4°C | ∞ | |

3.3. Take 3 µl aliquots after 8,10, 12 and 14 cycles at the last 15 seconds of elongation.
(Negative control - only at the end)

3.4. Prepare gel loading mix: UPW 7 µl + Loading dye 2 µl, distribute 9 µl to each tube.

3.5. Run on 1% agarose gel. Choose the minimal cycle for amplification.
If there is a problem with the gel check concentration with HS Denovix.

3.6. Do not proceed to production PCR in the same day.

4. Production PCR 2

4.1. Set up in UV cabinet 2X50 µl reactions:

| | X1 | X2 |
|---|---|---|
| DNA library | 10 µl | 20 µl |
| Kappa HiFi HS ready mix | 25 µl | 50 µl |
| Ilumina For | 2.5 µl | 5 µl |
| Ilumina Rev | 2.5 µl | 5 µl |
| UPW | 10 µl | 20 µl |
| Total | 50 µl | 100 µl |

4.2. Run PCR program according to calibration.

| Step | Temp | Time |
|---|---|---|
| 1 | 95 C° | 3 min |
| 2 | 98 C° | 20 sec |
| 3 | 65 C° | 30 sec |
| 5 | 72 C° | 45 sec |
| 6 | 72 C° | 3 min |
| 7 | 4°C | ∞ |

4.3. Purify the reaction with Ampure XP beads:

4.3.1. Combine two reaction to the same tube.

4.3.2. Add 1.7 volumes of Ampure beads (resuspend thoroughly) to the tube (170 µl).

4.3.3. Incubate for 10 min at RT. Place on MPS.

4.3.4. Wash twice with 80% ethanol: I) up-and-downs 3-4 times, II) incubate for 5 min on MPS.

4.3.5. Air-dry the beads.

4.3.6. Resuspend the beads in 25 µl Tris-HCl 10mM, incubate for 5 min, put on MPS.

4.3.7. Transfer the liquid to a new tube.

4.3.8. Check concentration with HS Denovix.

4.3.9. Run 1 µl on 1% Agarose gel.

### F. Oligo and primer sequences

| | |
|---|---|
| **Micro-CTALE oligos** | |
| BAC CH17-219E19 | chr1:30754221-30968290 |
| BAC-adapter-for | CCATCTCATCCCTGCGTGTCGACTACACTACTCGT |
| BAC-adapter-rev | /5Phos/CGAGTAGTGTTCAGCAAGGCACACAGGGGATAGG |
| BAC-bio-for | /5Biosg/CCATCTCATCCCTGCGTGTC |
| Bac-bio-rev | /5Biosg/CCTATCCCCTGTGTGCCTTG |
| BAC-rev | CCTATCCCCGTGCCTTG |
| Indexed primers WT | i5 barcode – GTACACCT i7 barcode – TCCTCATG |
| Indexed primers CTCF insert | i5 barcode – CAATGCGA i7 barcode – GACGAACT |
| Illumina uni oligo | ACACTCTTTCCCTACACGACGCTCTTCCGATCT |
| Illumina-for | AATGATACGGCGACCACCGAGAT |
| Illumina-rev | CAAGCAGAAGACGGCATACGA |
| **CRISPR/CTCF insert clone generation oligos** | |
| CRM_1A_F | GTTGTTTATGGAGCAGGTGAGG |
| CRM_1A_R | CTCAAGGTTTGAAGGAGGTGAC |
| CRM_2C_F | CCAGGTCACCATCCGGAC |
| CRM_2C_R | TGGAGACCTGTGGATAGG |
| g-ch1-313f | CACCGCAGGTCACCATCCGGACAAC |
| g-ch1-313r | AAACGTTGTCCGGATGGTGACCTGC |
| Inserted sequence (Chr2:43,056,093-43,056,234) | TCCTTGTGCAGTTCCCCCAGGGCTAATGGGGTTTGGTTGCCCACCTTTGCTGTGTCTCTT CGTGGCGCCAGCAGGTGGCAGCATTGCCTTTGCATGTGCCGGCTCCACCTCTGGCCTCT CCCAGACATGCAGCGTGTAGGG |

### G. Reagents and equipment

| Reagent | Manufacturer | Catalog Number |
|---|---|---|
| 37% Formaldehyde | Sigma Aldrich | 252549 |
| AMPure XP beads | Beckman Coulter | A63881 |
| CaCl2 | Spectrum Chemical | c1096 |
| Cot-1 DNA | InvivoGen | 15279011 |
| Denhardt's solution (50X) | Sigma-Aldrich | D2532-5ML |
| Digitonin | Sigma-Aldrich | 300410 |
| DNA LoBind tubes, 1.5 ml | Axygen | 81876MCT175LC |
| DNA polymerase I, large (Klenow) fragment | NEB | M0210S |
| ATP (part of dNTP set) | NEB | N0446S |
| dsDNA High Sensitivity Assay | DeNovix | IM-DSDNA-H2 |
| Dynabeads MyOne Streptavidin C1 | Invitrogen | 650.01 |
| Dynal MPC-S Magnetic Particle Concentrator (MPS) | Invitrogen | 120.20D |
| EcoRI | NEB | R0101S |
| Fetal bovine serum | Biological Industries | 04-121-1A |
| Halt Protease Inhibitor Cocktail (100X) | Thermo Fisher | 78429 |
| HEPES | Sigma-Aldrich | H3375 |
| HindIII | NEB | R3104M |
| Iscove's Modified Dulbecco's Medium (IMDM) | Biological Industries | 01-058-1A |
| jetOPTIMUS reagent | Polyplus | 101000025 |
| KAPA HiFi HotStart ReadyMix | Roche | KK2601 |
| MgCl2 | Sigma-Aldrich | M3634 |
| Micrococcal nuclease (MNase) | NEB | M0247S |
| NEBNext Multiplex Oligos for Illumina | NEB | E6440S |
| NEBNext Ultra II DNA Library Prep Kit | NEB | E7645S |
| Phenol–chloroform | Bio-Lab | 1697234400 |
| PMSF | Thermo Fisher | 36978 |
| Proteinase K | NEB | P810S |
| RNase A | Invitrogen | AM2270 |
| SDS (10%) Solution | Biological Industry | 01-890-1B |
| Sheared salmon sperm DNA | Sigma | D-7656 |
| T4 DNA ligase 400 U/µl | NEB | M0202L |
| T4 polynucleotide kinase (PNK) | NEB | M0201 |
| Trypsin-EDTA | Sigma-Aldrich | T4049 |
| Covaris microTUBE | Covaris | 520045 |
| Cells and Tissue DNA Isolation Kit | Norgen | 53100 |
| T4 DNA polymerase | NEB | M0203 |
| DpnII | NEB | R0543 |
| Biotin-14-dATP | Jena BioScience | NU-835-Bio14-S |
| USER Enzyme | NEB | E7645 |
